# Co-deletion of *ATAD1* with *PTEN* primes cells for BIM-mediated apoptosis

**DOI:** 10.1101/2021.07.01.450781

**Authors:** Jacob M. Winter, Heidi L. Fresenius, Heather R. Keys, Corey N. Cunningham, Jeremy Ryan, Deepika Sirohi, Jordan A. Berg, Sheryl R. Tripp, Paige Barta, Neeraj Agarwal, Anthony Letai, David M. Sabatini, Matthew L. Wohlever, Jared Rutter

**Author notes:** Correspondence should be addressed to J.R.

## Abstract

*PTEN* is a potent tumor suppressor gene that is frequently mutated or deleted in human cancers. Such deletions often include portions of the 10q23 locus beyond the bounds of *PTEN* itself, in many cases resulting in the disruption of additional genes. Coincidental loss of *PTEN*-adjacent genes might impose vulnerabilities that could either affect patient outcome basally or be exploited therapeutically. Here we describe how the loss of *ATAD1*, which is adjacent to and frequently co-deleted with *PTEN*, predisposes cancer cells to apoptosis and correlates with improved survival in cancer patients. ATAD1 directly and specifically extracts the pro-apoptotic BIM protein from mitochondria to inactivate it. Cells lacking ATAD1 are hypersensitive to clinically used proteasome inhibitors, which increase BIM and trigger apoptosis. Thus, we demonstrate that mitochondrial protein quality control interfaces with cell death in a clinically actionable manner.

## Main Text

The tumor suppressor gene *PTEN* is one of the most commonly deleted genes in human cancers, being deleted in more than 33% of metastatic prostate tumors and nearly 10% of glioblastoma multiforme and melanoma ^1^. Such deletions are imprecise and typically include *PTEN* as well as neighboring loci on chromosome 10q23. Only 40 kb upstream of *PTEN* is *ATAD1*, which encodes a AAA+ ATPase involved in protein homeostasis on the outer mitochondrial membrane (OMM) ^2–6^. *ATAD1* is essential for life in mammals and it has been conserved over the 1 billion years of evolution separating yeast and humans ^6,7^. Here we describe how co-deletion of *ATAD1* with *PTEN* sensitizes cells for apoptosis.

Because the *PTEN* and *ATAD1* genes are adjacent on human Chr10q23.31 (Fig 1A) ^8^, we assessed whether *ATAD1* is co-deleted with *PTEN* using immunohistochemistry on prostate adenocarcinoma (PrAd) tumors ^9^. We analyzed tumors that were *PTEN*-null by clinical sequencing (FoundationOne), along with *PTEN*-wild-type controls. ATAD1 protein was undetectable in more than half of the *PTEN*-null tumors analyzed (21/37), but was present in all 15 PTEN-wild-type control tumors (Fig 1B,C, Fig S1A). Analysis of genomic data from The Cancer Genome Atlas corroborated these protein-level findings, as the majority of tumors harboring deep deletions in *PTEN* also had deep deletions in *ATAD1* (Fig S1B). Importantly, *ATAD1* is almost never deleted in the absence of *PTEN* deletion (Fig S1B), nor does it feature recurrent inactivating point mutations (Fig S1C), which argues that ATAD1 is not a tumor suppressor itself. Therefore, we hypothesize that *ATAD1* deletion is simply a “hitchhiker” with the oncogenic driver of *PTEN* deletion. Nonetheless, *ATAD1* is deleted at a high frequency across many tumor types, including in more than 25% of PrAd, 11% of melanoma, and 7-8% of glioblastoma and lung squamous cell carcinoma (Fig 1D). Given the established role of ATAD1 in mitochondrial homeostasis, we hypothesized that a hitchhiker deletion of *ATAD1* would confer unique vulnerabilities on tumors ^10–13^.

**Figure 1:**
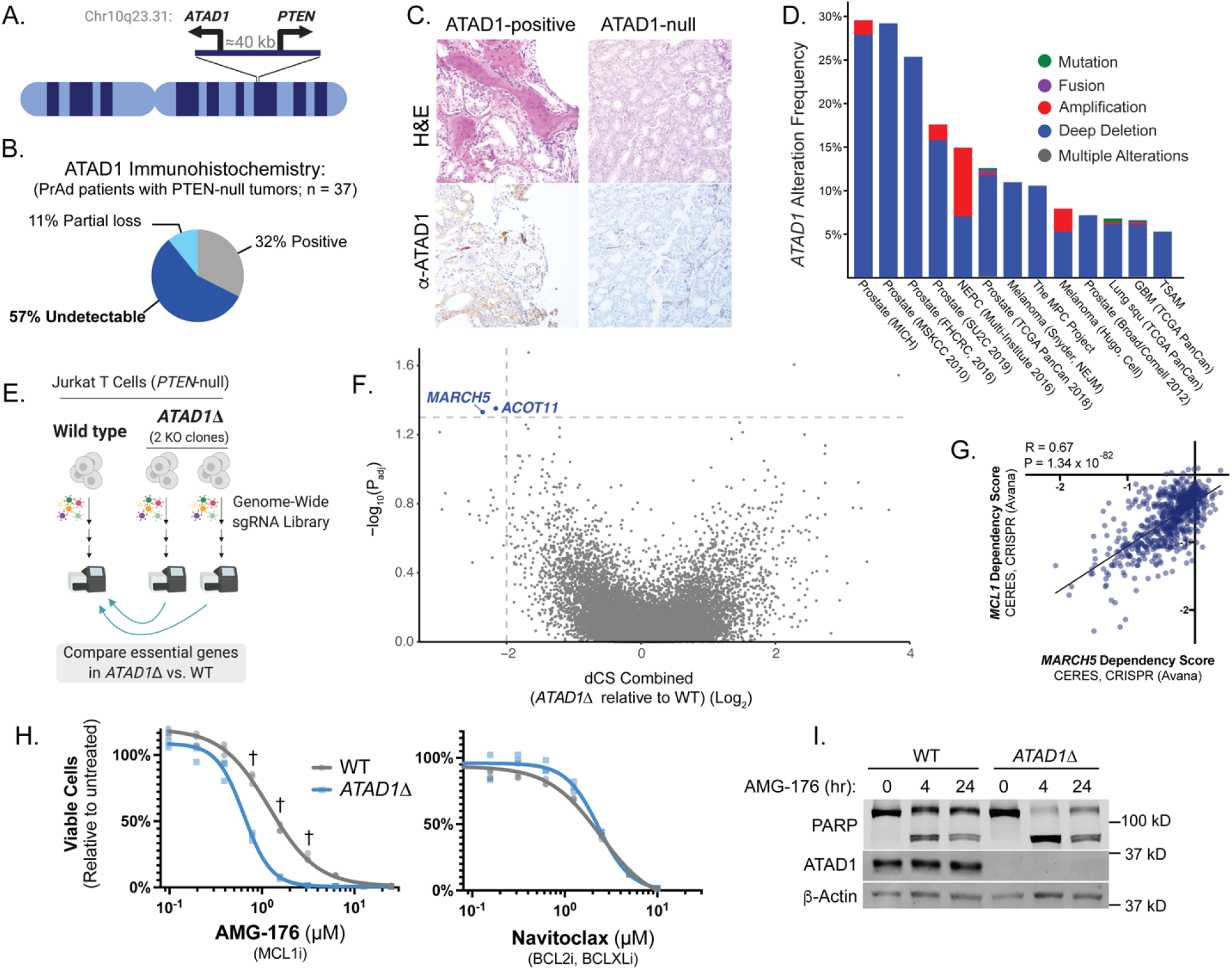
*ATAD1* is co-deleted with *PTEN* and its loss increases dependency on *MARCH5*/*MCL1*. (A) Schematic depicting the juxtaposition of ATAD1 and PTEN on human Chr10q23 (B) Summary of immunohistochemistry data from anti-ATAD1 staining of samples from prostate cancer patients with PTEN-null tumors (C) Representative histology from patients as described in B (D) Frequency of genetic alterations at *ATAD1* across various types of human cancer (E) Schematic depicting CRISPR screening strategy (F) Volcano plot of CRISPR screening data. X-axis depicts combined differential CRISPR Score (dCS) from the two ATAD1Δ clones relative to WT while the y-axis depicts adjusted P-value. Dashed lines demarcate P < 0.05 and dCS(Log2) < -2. (G) Gene essentiality data from Depmap revealing that *MARCH5* is co-essential with *MCL1* (H) Viability of Jurkat cells treated with AMG-176 (MCL1 inhibitor) or navitoclax (BCL2, BCLXL inhibitor) for 24 hours. N = 3 biological replicates representative of at least 2 independent experiments. (I) Western blot of whole cell lysates from Jurkat cells treated with 1 μM AMG176; Cleavage of PARP indicates caspase activity and is a measure of apoptosis. Representative of 3 independent experiments

To discover such vulnerabilities, we conducted genome-wide CRISPR-KO screens to identify genes that are selectively essential in *ATAD1*Δ cells. We generated two clonal *ATAD1*Δ lines in Jurkat cells, which are *PTEN*-null, using transient expression of Cas9 and two sgRNAs targeting distinct exons of *ATAD1* (Fig S2A,B). *ATAD1* deletion did not affect basal proliferation rate (Fig S2C). We conducted parallel screens on the wild-type (WT) Jurkat parental cell line and each of two *ATAD1*Δ clonal cell lines, the comparison of which enabled us to minimize idiosyncrasies inherent to clonal cell lines (Fig 1E). Differential CRISPR Scores (dCS) represent the difference between WT and *ATAD1*Δ in fold-change of abundance of sgRNAs targeting a given gene. By inference, genes with negative scores are more important for cell fitness in cells lacking *ATAD1*, compared to wild-type cells. As expected, dCS values for the two clonal cell lines significantly correlated (Fig. S2D). P values were multiplied to prioritize genes that scored as hits in both of the two *ATAD1*Δ clones ^14^. Two genes, *ACOT11* and *MARCH5*, met our statistical criteria of a combined dCS value of < -2 and P_adj_ < 0.05 (Fig 1F, Fig S2E,F). *ACOT11* encodes an acyl-CoA thioesterase that localizes to mitochondria and the cytoplasm and hydrolyzes long-chain fatty acyl-CoAs ^15,16^. *MARCH5* encodes a ubiquitin E3 ligase localized to the OMM that targets a number of transmembrane proteins for degradation ^17–19^. We chose to focus on the genetic interaction between *ATAD1* and *MARCH5* given the potential functional relationship between these two OMM-localized proteins involved in protein homeostasis.

MARCH5 was recently shown to promote the degradation of certain proteins of the BCL2 family, which function at the OMM to positively or negatively regulate the initiation of apoptosis ^20–23^. The critical step in the regulation of apoptosis is the permeabilization of the OMM by BAX/BAK, which causes mitochondrial proteins such as cytochrome c to diffuse into the cytoplasm and activate the caspase cascade. Anti-apoptotic proteins, including BCL2, BCL-XL, and MCL1, bind to and inhibit BAX and BAK ^22^. In response to a pro-death signal, the BH3-only proteins (including BIM, BID, NOXA, BAD, PUMA, etc.) trigger apoptosis by inactivating the anti-apoptotic proteins or directly activating BAX/BAK ^24^. Deletion of *MARCH5* causes MCL1, NOXA, and BIM to accumulate on the OMM, and increases sensitivity to BH3 mimetic drugs, which sequester anti-apoptotic BCL2 family members to trigger apoptosis ^25–28^.

Moreover, evidence from genetic screens supports the putative role of MARCH5 in the regulation of apoptosis by MCL1 ^28,29^. For instance, dependency on *MARCH5* for fitness is the best predictor of *MCL1* dependency (and vice versa) across the entire genome in the 859 cell lines used in the DepMap project ^30^. *MCL1* itself was not covered in our screen due to low abundance of sgRNAs at the initial time point. The three sgRNAs targeting *MCL1* that were adequately represented all show the expected trend of increased depletion in *ATAD1*Δ cells relative to that seen in wild type cells (Fig S2G).

Since dependence on *MARCH5* predicts dependence on *MCL1*, we tested the effect of *ATAD1* deletion on sensitivity to a MCL1-specific inhibitor, AMG176 ^31^. As predicted by our CRISPR screens, *ATAD1* loss sensitized to apoptosis induced by AMG176, but not by navitoclax (BCL2/BCLXL inhibitor), indicating that ATAD1 suppresses apoptosis in a mechanism-specific manner (Fig 1H,I) ^31^. In sum, through unbiased genetic screens we identify that ATAD1 loss increases dependency on the MARCH5/MCL1 anti-apoptotic axis.

The synthetic lethality of *ATAD1* and *MARCH5* in our screen suggested that they likely act in parallel pathways. Indeed, like MARCH5, deletion of *ATAD1* increased the abundance of both MCL1 and BIM, while having no effect on the abundance of FIS1, a tail-anchored protein on the OMM that is not part of the BCL2 family (Fig 2A,B). Thus, similar to MARCH5, ATAD1 regulates the abundance of members the BCL2 family and affects the sensitivity of cells to pro-apoptotic stimuli.

**Figure 2:**
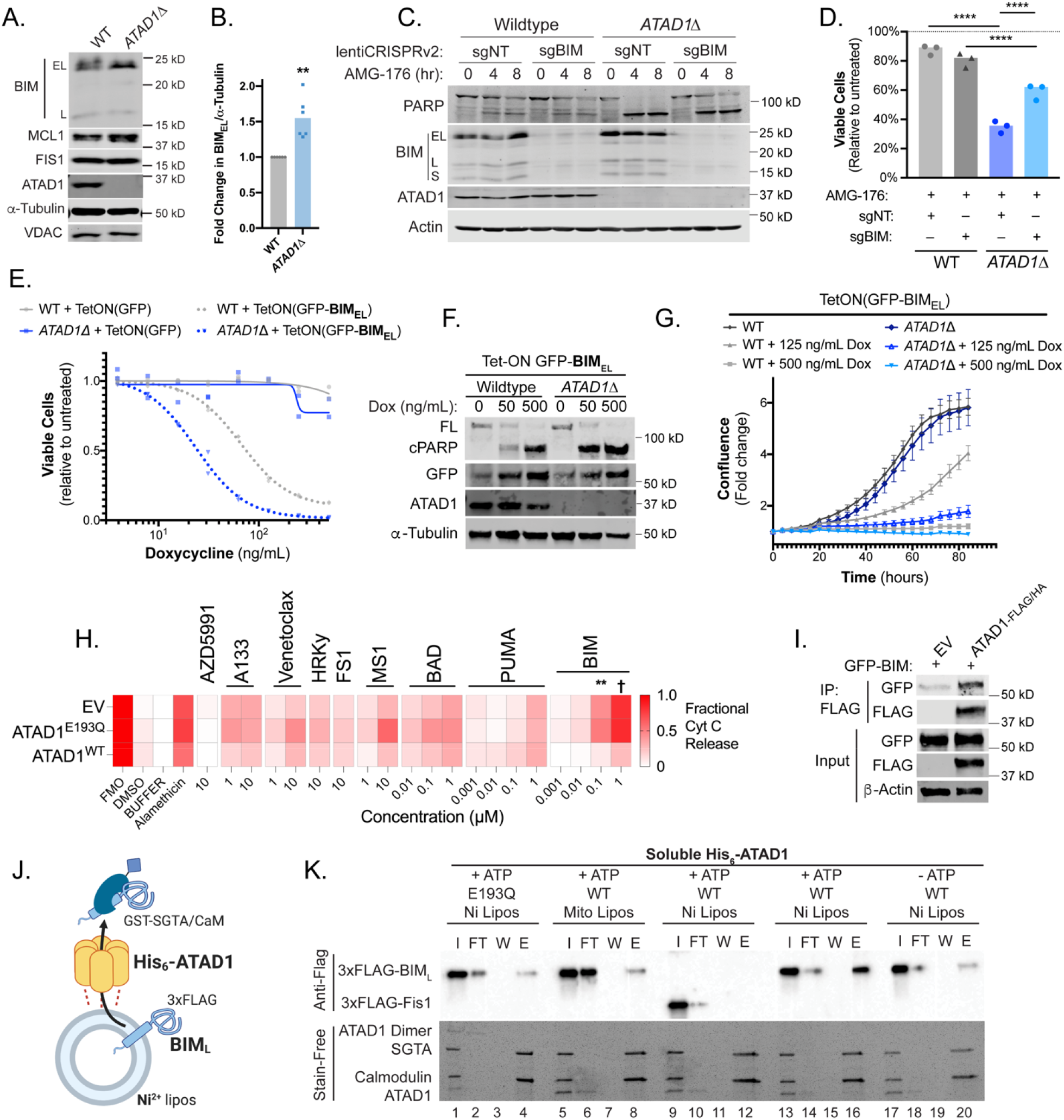
ATAD1 inactivates BIM by directly extracting it from membranes. (A) Western blot of whole-cell lysates from Jurkat cells (B) Quantification of BIM_EL_ levels normalized to alpha-tubulin levels in WT and *ATAD1*Δ cell lines; n = 6 independent experiments (C) Western blot of WT or *ATAD1*Δ cells stably expressing Cas9 and the indicated sgRNA, treated with 1 μM AMG-176 (D) Viability of Jurkat cells treated with 1 μM AMG-176 and expressing the indicated sgRNAs along with Cas9; n = 3 biological replicates (E) Viability of Jurkat cells transduced with tetracycline-inducible GFP-BIM_EL_ or GFP only and treated with varying concentrations of doxycycline for 48 hr; n = 2 biological replicates representative of 3 independent experiments (F) Western blot of lysates from WT or *ATAD1*Δ cells with Tet-ON(GFP-BIM_EL_) treated with indicated concentrations of docycycline for 24 hr; n = 3 independent experiments (G) Fold change in confluence measured by Incucyte software, of cell lines described in F, treated with indicated concentrations of doxycycline; mean ± SEM from n = 3 biological replicates, representative of 3 independent experiments (H) BH3 profiling data on H4 glioma cells (Del10q23) transduced with the indicated constructs (rows). A133 indicates A1331852. FMO: Fluorescence Minus One FACS control; Mean of n = 3 biological replicates is shown; † indicates P < 0.001; (I) Western blot of co-immunoprecipitation from H4 cells transduced with EV or ATAD1-FLAG/HA and transfected with GFP-BIM_EL_ in the presence of zVAD-FMK. Representative of 2 independent experiments (J) Schematic of reconstituted proteoliposome extraction assay (K) Extraction assay using His-ATAD1 and 3xFLAG-BIM_L_ (lanes 1-4, 9-20) or the negative control TA protein, 3xFLAG-Fis1 (lanes 5-8); GST-tagged SGTA and calmodulin (CaM) are included as chaperones to catch extracted TA protein. “I” = Input, “FT” = flow-through, “W” = final wash, “E” = elution. Eluted fractions represent TA proteins extracted by ATAD1 and bound by GST-tagged chaperones; compare elution “E” to input “I”. † denotes P < 0.0001

We hypothesized that the increased abundance or activity of BIM was responsible for the increased apoptosis of *ATAD1*Δ cells in response to MCL1 inhibition. Indeed, CRISPR-mediated deletion of *BCL2L11* (encoding BIM; “sgBIM”) partially rescued the AMG176 inhibitor sensitivity of *ATAD1*Δ cells as measured by PARP cleavage and cell viability assays (Fig 2C,D). Having demonstrated its partial necessity, we next asked if BIM was sufficient to trigger apoptosis in *ATAD1*Δ cells. BIM exists in three main isoforms, BIM_EL_ (“Extra-Long”, the predominant form), BIM_L_ (“Long”), and BIM_S_ (“Short”), which share key structural features including a C-terminal membrane anchor ^32–34^. We generated Jurkat cell lines expressing a tetracycline-inducible GFP-BIM_EL_ fusion. *ATAD1*Δ cells were hypersensitive to ectopic BIM_EL_, as measured by cell viability assays (Fig 2E), cleaved PARP immunoblot (Fig 2F), and live cell imaging using an Incucyte platform (Fig 2G). Importantly, maximal expression of GFP-BIM_EL_ using 500 ng/mL doxycycline killed cells regardless of the presence or absence of ATAD1. Thus, endogenous ATAD1 protects against BIM_EL_ but ATAD1 can be overwhelmed with sufficient levels of BIM. Together, these results demonstrate that BIM is sufficient and partially necessary for the apoptosis sensitivity of cells lacking ATAD1.

That ATAD1 suppressed BIM-induced cell death suggested that cells with Chr10q23 deletion, which endogenously lack ATAD1, might be “primed” for apoptosis. To address this possibility, we used the gold-standard assay of apoptotic priming known as BH3 profiling ^35^. We treated permeabilized cells with the isolated BH3 domain from different pro-apoptotic BH3-only proteins, and mitochondrial outer membrane permeability was measured by detecting cytochrome c release from mitochondria. We profiled H4 glioma cells (Del(10q23)) in which we had stably re-expressed ATAD1, a catalytically inactive mutant ATAD1^E193Q^, or the empty vector (EV). In the ATAD1-null cells transduced with EV or ATAD1^E193Q^, BIM peptide—even at nanomolar concentrations—caused nearly complete release of cytochrome c. Re-expression of wild-type ATAD1 protected cells from BIM-mediated cytochrome c release (Fig 2H). In contrast, ATAD1 status did not affect cytochrome c release induced by other BH3 peptides, suggesting that BIM in particular can exploit the absence of ATAD1. Alamethicin, a pore-forming peptide that permeabilizes the OMM independently of BAX/BAK and BH3-only proteins, induced cytochrome c release equally regardless of ATAD1 status, which indicates that ATAD1 does not affect cytochrome c release downstream of pore formation on the OMM. Thus, ATAD1 specifically suppresses apoptotic priming that is sensitive to BIM peptide by exclusively acting upstream of BAX/BAK. To summarize, our biochemical and genetic data indicate that ATAD1 deficiency sensitizes to BIM, a BH3-only protein that potently induces apoptosis.

We hypothesized that BIM might be a direct substrate of the ATAD1 extractase, which could explain how ATAD1 represses BIM-induced apoptosis. BIM resembles known ATAD1/Msp1 substrates; it is tail-anchored, intrinsically disordered, has basic residues C-terminal to its transmembrane domain, and is degraded by the proteasome ^36–38^. Consistent with BIM being an ATAD1 substrate, GFP-BIM_EL_ co-immunoprecipitated with FLAG-tagged ATAD1 in H4 cells (Fig 2I). We further tested whether ATAD1 can directly extract BIM from a membrane using an in vitro system. We used BIM_L_ because it is more soluble than BIM_EL_ but shares the key structural features that would likely mediate ATAD1 recognition, including the tail-anchor and juxtamembrane regions ^32^. We were unable to purify active, full-length ATAD1. Instead, we swapped the N-terminal transmembrane domain with a His_6_ tag, which anchored His_6_-ATAD1 to liposomes doped with phospholipids containing Nickel-chelated headgroups (“Ni Lipos”, Fig 2J). In this extraction assay, tail-anchored (TA) proteins that are extracted from liposomes by ATAD1 are bound by soluble GST-tagged chaperones (SGTA and calmodulin), purified on a glutathione column, and detected by Western blot ^38^. We validated the Ni-His anchoring strategy using full-length or truncated yeast Msp1 and positive and negative control substrates (Fig S3A-C).

His_6_-ATAD1 directly and efficiently extracted 3xFLAG-BIM_L_ from liposomes in this assay (Fig 2K). As expected, this activity was ATP-dependent (lanes 17-20) and was abolished when we used the catalytically inactive mutant, ATAD1^E193Q^ (lanes 1-4). Swapping the Ni-chelating lipids for lipids typical of the OMM (“Mito Lipos,” lanes 5-8, which cannot anchor His_6_-ATAD1) prevented extraction of BIM, demonstrating that ATAD1 requires membrane anchoring for its extractase activity. Importantly, ATAD1 did not extract Fis1, another TA protein (lanes 9-12). We conclude from these results that ATAD1 specifically extracts BIM, but not all TA proteins, from lipid membranes.

These data raise the question of what happens to BIM within the cell after it has been extracted by ATAD1. We transduced SW1088 cells, which are a large and flat Del(10q23) cell line suitable for imaging, with either EV or ATAD1-FLAG. These cells were further transduced with TetON(GFP-BIM_EL_ΔBH3), in which four point mutations in the BH3 domain of BIM neutralize its pro-apoptotic activity, which permits live cell imaging. We assessed GFP-BIM_EL_ΔBH3 localization using live cell confocal microscopy in the absence and presence of ATAD1, using MitoTracker Red (MT-Red) to label mitochondria. ATAD1 altered the localization of BIM under basal conditions, generating GFP-positive puncta that did not colocalize with mitochondria (Fig 3A). Since BIM is regulated by proteasomal degradation, we additionally treated cells with bortezomib (BTZ), a proteasome inhibitor. Treatment with BTZ exacerbated this phenotype and resulted in larger, brighter GFP+ puncta only in ATAD1 expressing cells (Fig 3B, Fig S4). Thus, ATAD1 shifts the localization of BIM from mitochondria to cytoplasmic puncta, consistent with BIM being a substrate for mitochondrial extraction.

**Figure 3:**
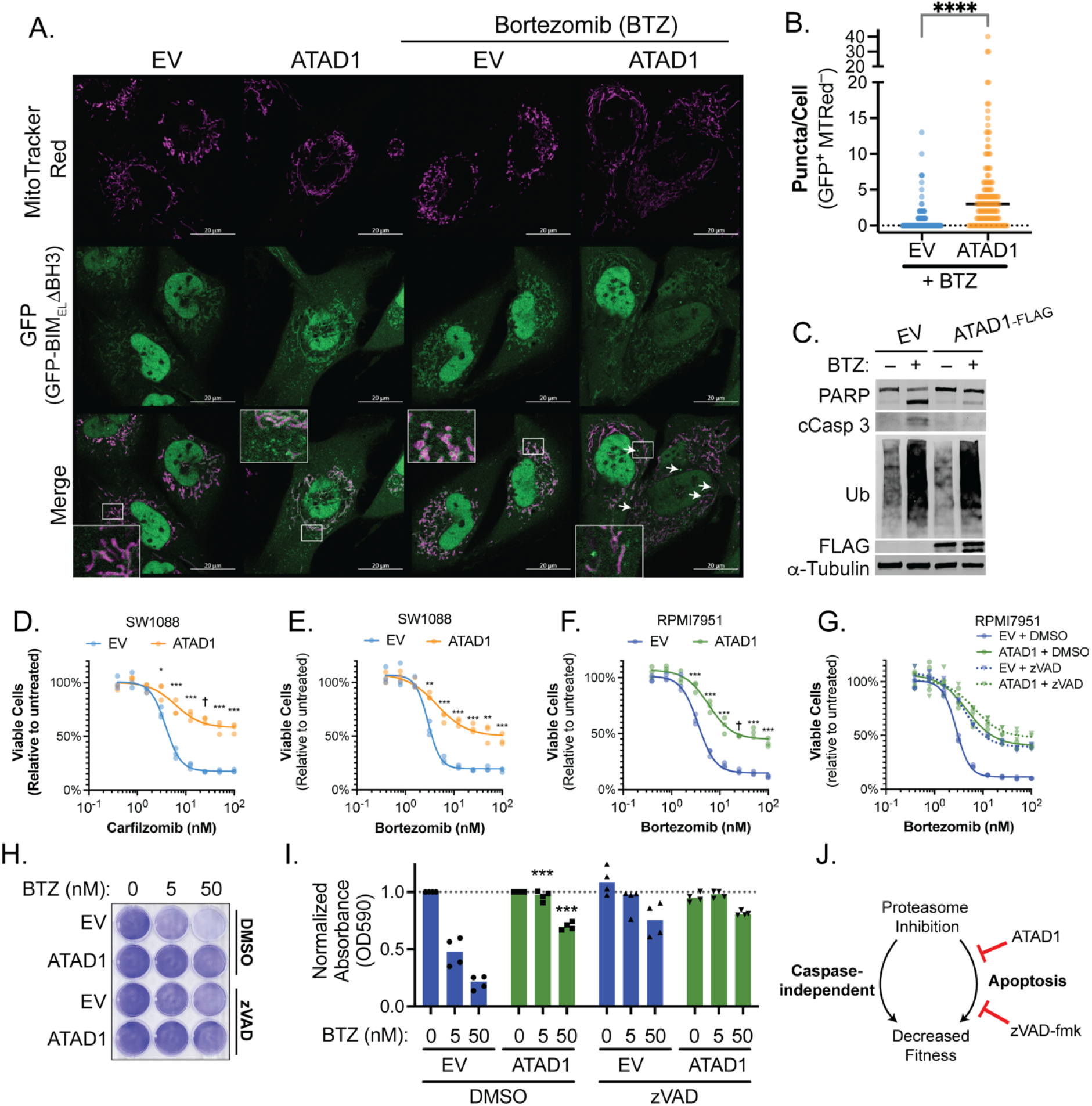
ATAD1 protects cells from apoptosis induced by proteasome dysfunction. (A) Confocal microscopy of live cells transduced with EV or ATAD1-FLAG, plus TetON(GFP-BIM_EL_ΔBH3) and treated with 100 ng/mL doxycycline for 24 hr. Bortezomib treatment was at a concentration of 100 nM for 2 hr. Mitochondria were visualized with MitoTracker Red. Images are representative of at least 3 independent experiments. Arrows indicate GFP^+^ MTRed^−^ puncta. Scale bar = 20 μm (B) Quantification of GFP^+^ MTRed^−^ puncta in BTZ-treated cells, as shown in (A); n = 132 (EV) or 127 cells (ATAD1) compiled from 3 independent experiments. (C) Western blot of PARP cleavage and poly-ubiquitin accumulation upon treatment with bortezomib (BTZ) at 10 nM for 8 hr; representative blots from n = 3 independent experiments, quantified in (C). (D) Viability of SW1088 cells transduced with EV or ATAD1-FLAG and treated with various concentrations of carfilzomib or bortezomib (E) for 24 hr. (F) Viability of RPMI7951 cells treated with bortezomib for 16 hr. (G) Viability of RPMI7951 cells co-treated with bortezomib and DMSO (vehicle) or zVAD-FMK (20 μM) for 16 hr. n = 3 biological replicates representative of 2 independent experiments. (H) Crystal violet staining of RPMI7951 cells treated with indicated compounds and concentrations for 16 hr and quantified in (I); n = 4 independent experiments (J) Schematic depicting how ATAD1 protects cells from apoptosis, but not caspase-independent cell death, in response to proteasome inhibition.

BIM accumulates upon dysfunction of the ubiquitin proteasome system induced by stressors such as loss of MARCH5 or treatment with proteasome inhibitors (Fig S5A,B)^39^. We hypothesized that cell death caused by blocking BIM degradation using proteasome inhibitors in Del(10q23) cells could synergize with the absence of ATAD1. Bortezomib (BTZ) robustly and rapidly induced apoptosis in Del(10q23) cells (RPMI7951 melanoma) transduced with EV, but not in cells in which we re-expressed ATAD1 (Fig 3C, S5C). Polyubiquitinated proteins accumulated to the same extent in the presence or absence of ATAD1, indicating that ATAD1 affects how the cell responds to proteotoxic stress, rather than minimizing the proteotoxic insult itself. Further, re-expression of ATAD1 mitigated cell death induced by multiple, structurally distinct proteasome inhibitors in various Del(10q23) cell lines (Fig 3D-F; Fig S5D-J). There are no available Del(10q23) prostate cancer cell lines, but PC3 cells are PTEN-null and *ATAD1*-hemizygous (Fig S5K). Deletion of the remaining allele of *ATAD1* in PC3 cells sensitized to BTZ, while overexpression of ATAD1^WT^ – but not ATAD1^E193Q^ – increased resistance to BTZ (Fig S5L-N). Thus, ATAD1 becomes essential for viability in cells subjected to ubiquitin proteasome system dysfunction.

Apoptotic cell death is only one of many mechanisms underlying proteasome inhibitor toxicity in cells ^40–42^. We next asked whether ATAD1 affects proteasome inhibitor sensitivity via some apoptosis-independent pathway, in which case ATAD1 re-expression and caspase inhibition (which blocks apoptosis) would have an additive effect in mitigating proteasome inhibitor toxicity. Strikingly, ATAD1 re-expression completely phenocopied treatment with the caspase inhibitor zVAD-fmk in Del(10q23) cells treated with BTZ (Fig 3G–I). Moreover, there was no additive effect of zVAD-fmk in ATAD1 re-expressing cells, suggesting that ATAD1 and zVAD-fmk act in the same pathway to prevent the death of cells subjected to proteasome inhibition (Fig 3J). Together, these results indicate that the protective effects of ATAD1 during proteasome inhibition can be explained exclusively by limiting apoptosis.

Given our findings that ATAD1 regulates BIM, we examined whether ATAD1 status affects tumor growth in vivo. SW1088 is a Del(10q23) glioma cell line that is non-tumorigenic in SCID mice ^43,44^. As expected, SW1088 cells transduced with EV failed to form tumors in 17 out of 17 NOD/SCID mice. However, SW1088 cells transduced with ATAD1 grew palpable tumors in 17 out of 18 mice (Fig 4A,B). Thus, ATAD1 serves a pro-survival role in PTEN-null tumor cells in vivo, consistent with our findings that ATAD1 extracts BIM for degradation and protects against proteotoxic stress-induced apoptosis.

**Figure 4:**
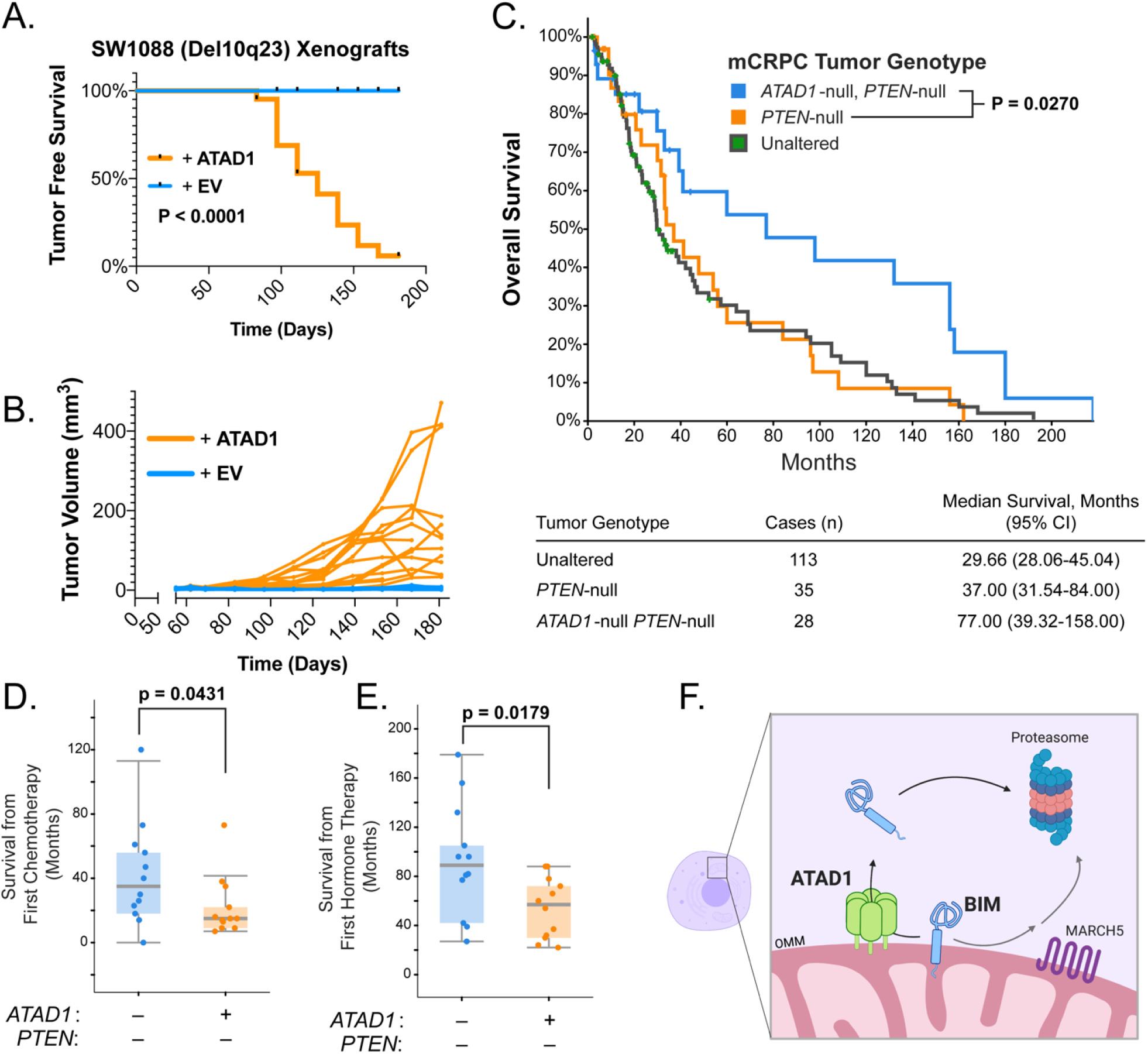
ATAD1 exerts a pro-tumor effect in murine xenografts and human prostate cancer patients. (A) Kaplan-Meier curve of tumor-free survival of mice injected subcutaneously with SW1088 cells transduced with EV (n = 17) or ATAD1-FLAG (n = 18). (B) Tumor volume measurements for mice described in (A) (C) Kaplan-Meier curve of overall survival from patients with metastatic castrate-resistant prostate cancer, stratified based on status at the ATAD1 and PTEN loci, with accompanying table below (D) Overall survival after initiating chemotherapy for a subset of mCRPC patients shown in (C); (E) Overall survival after initiating hormone therapy for a subset of mCRPC patients shown in (C); (F) Model of the relationship between ATAD1 and BIM

We next examined clinical outcomes in patients with tumors that were *PTEN*-null, both *PTEN*-null and *ATAD1*-null, or neither (“unaltered” at these loci). We queried two studies of patients with metastatic, castration-resistant prostate cancer (mCRPC), since *ATAD1* loss is prevalent in this tumor type^45,46^. The median overall survival of mCRPC patients with *PTEN*-null *ATAD1*-null tumors (77 months) was more than double that of patients with only *PTEN*-null or “unaltered” tumors (37 months, Fig 4C). In a subset of patients, we were able to additionally assess overall survival after initial hormone therapy or chemotherapy. Patients whose tumors lacked both *ATAD1* and *PTEN* had longer survival post-therapy than did the control patients whose tumors lacked only *PTEN* (Fig 4D,E). These clinical findings are consistent with our observation that ATAD1 suppresses apoptosis in cultured cells. Altogether, our data indicate that ATAD1 exerts a pro-survival effect in tumor cells (with ATAD1 deficiency decreasing tumor fitness) in both murine xenografts and human mCRPC patients.

It might appear counterintuitive that ATAD1 protects against apoptosis despite Chr10q23 deletion occurring so frequently in cancer. This is likely explained by the strong selection for deletion of *PTEN*, which virtually always co-occurs with deletion of *ATAD1*. PTEN loss activates AKT, which has strong pro-survival effects, including decreasing transcription of *BCL2L11* (BIM) by inhibiting FOXO3A, and inactivating BAD by direct phosphorylation ^47,48^. Thus, co-deletion of *PTEN* may buffer some of the apoptotic priming induced by *ATAD1* loss.

Neutral or detrimental alleles that act as genetic “hitchhikers” when they are physically linked to advantageous alleles are a well-described phenomenon in evolutionary biology ^49^. Additionally, genetic lesions that prime cells for apoptosis are not always selected against in cancer. On the contrary, potent oncogenes (including MYC paralogs) are strongly selected for despite priming for apoptosis ^50,51^. We cannot rule out the possibility that ATAD1 loss could be beneficial under some circumstances, but our data overwhelmingly support its role as a pro-survival factor.

The effect of ATAD1 on apoptosis likely extends beyond *PTEN*-null cancers into a broader physiological context. ATAD1 has not been explicitly linked to apoptosis in the literature, but it was originally discovered in mammals via a genetic screen for factors that prevent neuronal cell death in response to oxygen and glucose deprivation ^52^. Further, *Atad1* phenocopies *Bcl2* and *Bcl2l1* in the middle cerebral artery occlusion mouse model of ischemic stroke: deletion of *Atad1* or *Bcl2* exacerbates, and overexpression of *Atad1, Bcl2*, or *Bcl2l1* decreases neuronal cell death ^53–56^. Thus, we believe that antagonism of BIM likely contributes to the neuroprotective function of ATAD1 in vivo ^6^.

The delicate balance of pro and anti-apoptotic BCL2 family proteins constantly threatens cell survival. How these proteins are activated has been intensely studied, but less is known about how they are neutralized. Here we report that ATAD1 directly and specifically extracts OMM-localized BIM, leading to its inactivation (Fig 4F). Dysfunction of the ubiquitin proteasome system causes BIM to accumulate, and ATAD1 is essential for cell viability under such stressful conditions. That ATAD1 loss sensitizes to proteasome dysfunction could have therapeutic implications for thousands of cancer patients with Del10q23 tumors, given that drugs targeting the proteasome approved for the treatment of cancer. Thus, defining the mechanisms of ATAD1 regulation and function will reshape our understanding of apoptotic susceptibility, but it also has the potential for profound clinical impact for patients with *PTEN*/*ATAD1* co-deleted tumors.

## Supporting information

sgrna.counts

Gene.level.ATAD1.screen

mCRPC_patient_OS_data

## Acknowledgments

The authors acknowledge University of Utah Core Facilities, particularly James Marvin, PhD and the Flow Cytometry Core, David Lum, PhD and the Preclinical Research Resource, and the DNA/Peptide Synthesis Core. We thank Keren Hilgendorf for equipment. We thank Derick Torres for technical lab support. We thank Florian Muller for helpful conversations and insight. We thank members of the Rutter lab for helpful discussions and comments on the manuscript.

## Author Contributions

Conceptualization: JMW, JR; Methodology: JMW, HF, HK, CNC, J.Ryan, DS, TS, JAB, PB, MW; Formal analysis: JMW, HK, JAB; Investigation: JMW, HF, HK, CNC, J.Ryan, DS, TS; Resources: NA, AL, DMS; Writing-Original Draft: JMW, JR; Writing-Review & Editing: All authors; Supervision: JR; Project Administration: NA, AL, DMS, MW, JR; Funding acquisition: JMW, MW, JR

## Materials and Methods

### Lead Contact

Further information and requests for resources and reagents should be directed to and will be fulfilled by the lead contact, Jared Rutter.

### Materials Availability

All unique/stable reagents generated within this study are available from the lead contact upon request without restriction.

### Data and Code Availability

The CRISPR screening data generated during this study are available in the supplemental materials.

## EXPERIMENTAL MODEL AND SUBJECT DETAILS

### *ATAD1* knockout cell lines

Jurkat E6.1 human T-ALL cells (ATCC TIB-152) were grown in RPMI1640 with 10% FBS and 100 U/mL Pen/Strep (ThermoFisher). Cells were electroporated] using Lonza SE Cell Line 4D-Nucleofector™ X Kit L according to manufacturer’s specifications and protocol optimized for Jurkat E6.1 cells. Px458-derived plasmids encoding sgRNA targeting *ATAD1* were transiently expressed in Jurkat cells via electroporation. Three days later, GFP+ cells were sorted (BD FACSAria) and plated as single cells in 96 well plates. Clonal cell lines were grown, harvested, and evaluated for *ATAD1* deletion via immunoblot using a knockout-validated monoclonal antibody (NeuroMab).

PC3 cells were transduced with lentivirus encoding LentiCRISPRv2-GFP (LCv2G) with non-targeting sgRNA (sgNT; which contains a 15 nt sgRNA sequence, 5’-GAGACGGACGTCTCT-3’, that does not precede any 5’-NGG-3’ sites in the human genome. This was determined by using GAGACGGACGTCTCTNGG as input for ncbi BLAST, and filtering results to 100% query coverage, 100% identity) or LCv2G with sgATAD1 (guide sequence #1). Three days after transduction, GFP+ cells were sorted (BD FACSAria) and maintained as a polyclonal population. Editing was confirmed by immunoblot as above.

### *ATAD1* Re-expression in Del10q23 cell lines

H4 and PC3 cells were transduced with retrovirus (PQCXIP transfer plasmid) encoding ATAD1 with C-terminal FLAG and HA tags. Two days after transduction, cells were selected with 1 µg/mL puromycin for 4 days. RPMI7951, HGC27, and SW1088 cells were transduced with lentivirus (pLenti-Blast transfer plasmid) encoding ATAD1-FLAG, and selected with 8 µg/mL blasticidin for 6 days. Cells were grown in media containing the selective antibiotic upon thawing stocks, but no experiments were conducted using media that contained selective antibiotics.

## METHOD DETAILS

### Immunohistochemistry

#### ATAD1 (using NeuroMab #75-157 mouse monoclonal antibody)

The ATAD1 immunohistochemical staining was performed on 4-micron thick sections of formalin-fixed, paraffin-embedded tissues. Sections were air-dried and then melted in a 60°C oven for 30 minutes. Slides were loaded onto the Leica Bond™ III automated staining instrument (Leica Biosystems, Buffalo Grove, IL.) and de-paraffinized with the Bond™ Dewax solution. The antigen retrieval performed was done with Bond™ Epitope Retrieval Buffer 2 (ER2, pH 8.0) for 20 minutes at 95°C. The ATAD1 primary antibody concentration of 1:400 was applied at an incubation time of 30 minutes at room temperature. Positive signal was visualized using the Bond™ Polymer Refine Detection kit-DAB, which is a goat anti-mouse/anti-rabbit secondary HRP/polymer detection system, utilizing DAB (3-3’ diaminobenzidine) as the chromogen. Tissue sections were counterstained with hematoxylin for 10 minutes. The slides were removed from the immunostainer and placed in a dH2O/DAWN ™ mixture. The sections were gently washed in a mixture of de-ionized water and DAWN™ solution to remove any unbound reagent. The slides were gently rinsed in deionized water until all of wash mixture was removed. The slides were de-hydrated in graded ethanols, cleared in xylene and then coverslipped.

#### PTEN (using rabbit anti-human monoclonal antibody): clone 138G6, catalog # 9559L, Cell Signaling, Danvers, MA)

The PTEN immunohistochemical staining was performed on 4-micron thick sections of formalin-fixed, paraffin-embedded tissues. Sections were air-dried and then melted in a 60°C oven for 30 minutes. Slides were loaded onto the Ventana BenchMark™ Ultra automated staining instrument (Ventana Medical Systems, Tucson, AZ), de-paraffinized with the EZ Prep solution. The antigen retrieval performed was done with a citrate buffer (pH 6.0) in a pressure cooker (BioCare Medical, Concord, CA) for 4 minutes at 100°C then cooled in hot buffer for 30 minutes. The PTEN primary antibody concentration of 1:50 was applied at an incubation time of 2 hours at room temperature. The Ventana Amplification kit was applied to increase the antibody signal. Positive signal was visualized using the UltraView DAB detection kit, which is a goat anti-mouse/anti-rabbit secondary HRP/polymer detection system, utilizing DAB (3-3’ diaminobenzidine) as the chromogen. Tissue sections were counterstained with hematoxylin for 16 minutes. The slides were removed from the immunostainer and placed in a dH2O/DAWN ™ mixture. The sections were gently washed in a mixture of de-ionized water and DAWN™ solution to remove any unbound reagent and coverslip oil applied by the automated instrument. The slides were gently rinsed in deionized water until all of wash mixture was removed. The slides were de-hydrated in graded ethanols, cleared in xylene and then coverslipped.

### Cell Culture

Jurkat cells were cultured in RPMI1640 with 10% FBS (Sigma) and 100 U/mL Pen/Strep. Cells were counted regularly and typically split at a concentration of approximately 1-1.5 × 10^6^ cells/mL, but always before reaching a concentration of 3×10^6^ cells/mL. Jurkat cells transduced with tet-inducible vectors were cultured in RPMI1640 with 10% “Tet System Approved FBS” (Takara) instead of standard FBS. Adherent cell lines were maintained in subconfluent cultures in the following media: RPMI1640 (PC3), DMEM (H4, SW1088), EMEM (RPMI7951 and HGC27), all with 10% FBS and 100 U/mL Pen/Strep. Cells were periodically tested for mycoplasma contamination using a MycoAlert kit and were negative.

### Cloning

All cloning was conducted via traditional PCR/restriction enzyme “cut and paste” methods and verified by Sanger sequencing.

#### ATAD1 constructs

Retroviral plasmids encoding ATAD1-FLAG/HA and ATAD1^E193Q^-FLAG/HA were published previously (Chen et al, 2014). Lentiviral vectors were made, using the pLenti-BLAST backbone, by PCR-amplifying the ATAD1 CDS from the above retroviral vectors, but truncating the construct by replacing the HA tag with a stop codon, and ligating between SalI and XbaI sites.

#### GFP-BIM Constructs

The pLVXTet-One vector was purchased from Takara. The coding sequence for EGFP was PCR-amplified and ligated into the MCS using AgeI/BamHI sites. A fusion of EGFP-BIM_EL_ was generated using SOEing PCR and ligated using AgeI/BamHI sites. EGFP-BIM_EL_ was also ligated into pEGFP-C3 for transient transfection.

### Cell Titer Glo Viability Assay

Viability was determined by Cell Titer Glo (Promega) according to the manufacturer’s recommendation, with some modifications. Cells were plated at a density of 5 × 10^3^ cells/well (adherent cell lines) or 2-4 × 10^4^ cells/well (Jurkat) in 100 µL in 96 well plates with white walls and clear bottoms (Corning #3610). Cell Titer Glo reagent was reconstituted, diluted 1:4 using sterile PBS, and stored at -20 °C in 10 mL aliquots. The outer wells of the 96 well plates were filled with media but not with cells, due to concerns of edge effects. Luminescence was measured using a Biotek Synergy Neo2 microplate reader. Luminescence values were normalized on each plate to untreated cells on the same plate, and expressed as percent.

### Incucyte

Jurkat cells stably transduced with TetON(GFP-BIM_EL_) were seeded at a density of 10^3^ cells per well in clear-bottom, black 96 well plates (Corning) with different concentrations of doxycycline. Cells were imaged with an Incucyte SX5 system and monitored by phase contrast microscopy, with 5 images taken per well, every 4 hours, for approximately 4 days. Confluence was normalized to t = 0 and is expressed as fold change. Three replicate wells were used for each condition, and three independent experiments were conducted.

### Crystal Violet Staining

Cells cultured in 12 or 6 well plates were washed twice with PBS then fixed with 4% paraformaldehyde (Sigma Aldrich) for 30 minutes at room temperature. Wells were washed with ddH_2_O three times, then stained with 0.1% (w/v) crystal violet solution in 20% methanol for 30 minutes at room temperature. Wells were again washed with ddH_2_O three times, inverted to dry, and plates were photographed against a white background using an iPhone X. For quantification, glacial acetic acid was added to each well to elute the dye, and plates were incubated at room temperature on a rotary shaker for 30 minutes. Absorbance was measured at 590 nm using a Biotek Synergy Neo2 microplate reader, and values were normalized to those from untreated cells of the same genotype on each plate.

### SDS-PAGE and Immunoblotting

Whole cell lysates were prepared by scraping cells directly into RIPA buffer (or adding RIPA buffer to Jurkat cell pellets) supplemented with protease and phosphatase inhibitors (Sigma Aldrich P8340, Roche Molecular 04906845001), incubated on ice for 30 minutes with vortexing every 10 min, and then spun at 16,000 g for 10 minutes at 4°C to remove insoluble material.

Supernatant was saved as lysate and concentrations were normalized for total protein content after measuring with a BCA assay (Thermo Scientific 23225). Samples were resolved by SDS-PAGE or Tris-glycine gels (Invitrogen XP04205BOX) and transferred to nitrocellulose or PVDF (extraction assay) membranes. Immunoblotting was performed using the indicated primary antibodies which are listed in the key resources table according to the manufacturers’ recommendations, and analyzed by Licor Odyssey or Azure C500 (extraction assay). Note that the detector for the Azure C500 has several columns of pixels which appear to be non-functional. This gives the appearance of thin vertical white lines in some images. This can be readily viewed in raw data files by over-adjusting the contrast.

### Co-Immunoprecipitation

H4 cells (expressing EV or ATAD1-FLAG/HA) were transfected with GFP-BIM_EL_ in pEGFP-C3 (GFP-BIM), 10 µg plasmid for 10 cm plate, in the presence of 20 µM zVAD-fmk. Transient expression proceeded overnight (approximately 16 hr). Cells were washed with cold PBS and lysed with HN buffer supplemented with protease inhibitor cocktail and 1% CHAPS (HNC buffer). Magnetic anti-FLAG beads (Sigma Aldrich) were equilibrated with HNC buffer and then mixed with lysate (after removing 10% volume as input). Bead-lysate mixtures were incubated on a rotator at 4 °C for 2-4 hr. Beads were washed 3x with HNC buffer, then heated at 65 °C in 30 µL 1X Laemmli buffer for 10 min.

### Cell counting

Cells were counted using BioRad TC20 cell counter. At least two samples were taken from a culture any time a count was to be made and the mean was recorded. For proliferation experiments with Jurkat cells, a hemocytometer was used.

### CRISPR Screen

Jurkat cells (wildtype parental, *ATAD1*Δ #1, and *ATAD1*Δ #2) were transduced by spinfection with a genome-wide lentiviral sgRNA library (Addgene #1000000100; Wang et al, *Science* 2015) that also encoded Cas9 and a puromycin resistance cassette. Transduction was optimized to achieve an approximate transduction efficiency of 30%, and cells were selected with puromycin (0.5 µg/mL) for three days, allowed to recover without puromycin for two days, then maintained in a lower dose of puromycin (0.2 µg/mL) for the duration of the screen. An initial sample of cells (8 × 10^7^) were collected and frozen at the endpoint of puromycin selection (six days post-transduction). Cells were then maintained in culture for 14 cumulative population doublings. Cells were passaged every two days and seeded into new flasks at a density of 2 × 10^5^ cells/mL. After 14 population doublings, representative samples were collected (8 × 10^7^ cells).

As described elsewhere (Adelmann et al, *Methods in Mol Biol*, 2019), cell pellets were processed using a QIAamp DNA Blood Maxiprep, sgRNA sequences were amplified by PCR, and amplicons were sequenced for 40 cycles by Illumina HiSeq NGS at the Whitehead Institute DNA Sequencing Core Facility.

Sequencing reads were aligned to the sgRNA library, given a pseudocount of 1, and the abundance of each sgRNA was calculated as described previously (Wang et al, *Science* 2015; Kanarek et al, *Nature* 2018). The counts from each sample were normalized for sequencing depth. sgRNAs with fewer than 50 reads, and genes with fewer than 4 sgRNAs, in the initial reference dataset were omitted from downstream analyses. The log_2_ fold change in abundance of each sgRNA between the final and initial reference populations was calculated and used to define a CRISPR score for each gene. The CRISPR score is the average log_2_ fold change in abundance of all sgRNAs targeting a given gene. The distribution of all sgRNAs targeting a given gene was tested against the entire sgRNA distribution using the Kolmogorov-Smirnov test, and p-values were adjusted using the Benjamini-Hochberg procedure (Benjamini and Hochberg, *J. R. Statist. Soc*. 1995). To achieve a direct comparison of gene essentiality in an *ATAD1*Δ clone to that in the WT control, we omitted sgRNAs that were not adequately represented (i.e. < 50 reads at the initial time point) in both groups. This step enables a paired analysis of sgRNA changes in abundance, and avoids including a given sgRNA that “scored” in one genetic background but whose effects cannot be assessed in another. Differential CRISPR Scores (dCS) for each *ATAD1*Δ clone were calculated for each gene as: CS_ATAD1_Δ – CS_ATAD1-WT_. The mean of the two dCS values were summed and are presented as “dCS Combined.” Genes that scored as “hits” in one clone but not the other were penalized in the analysis by combining the Q values (Benjamini-Hochberg-corrected P values) for the two clones via Fisher’s Method (Fisher, R. A., 1934; also known as “sum of logs” method) for meta-analysis using the R package “metap” (Dewey M, 2020). Data were analyzed and plotted using ggplot2 with R version 4 and RStudio version 1.1.442

### BH3-Profiling

BH3-profiling was conducted using a FACS based method to directly monitor Cytochrome C release/retention in cells, as described previously (Ryan and Letai, *Methods* 2013).

### Confocal Microscopy

SW1088 cells transduced with EV or ATAD1-FLAG and TetON(GFP-BIM_EL_ΔBH3) were seeded at a density of 3.5 × 10^4^ cells per dish, in 35 mm Fluorodish plates (World Precision Instruments). Approximately 16 hours later, media was removed and replaced with media containing 100 ng/mL doxycycline. After 24 hours, cells were treated with 20 nM MitoTracker Red for 15 min and then imaged on a Zeiss LSM 880 confocal laser scanning microscope for 45 minutes in 5% CO_2_ at 37°C. Imaging on the Zeiss LSM 880 confocal laser scanning microscope was performed with a Plan-Apochromat 63x/1.40 Oil DIC f/ELYR objective. Alternatively, cells were treated with 100 nM bortezomib for 90 min prior to imaging, were stained with MitoTracker Red as described above, and imaged for 30 minutes. Doxycycline concentrations were maintained throughout the staining and imaging process. All images were airyscan processed using the Zeiss Zen Desk software.

Microscopy was conducted by an investigator (C. Cunningham) who was blinded to genotype (EV vs. ATAD1) and treatment (–/+ bortezomib). Experiments were repeated for 3 independent replicates (both – and + bortezomib) and 2 additional replicates (– bortezomib only). At least 40 images (representing approximately 40-60 cells) were taken per condition, per replicate. GFP-positive, MitoTracker Red-negative puncta were counted using FIJI with the multi-point tool and were graphed using GraphPad Prism 9 for MacOS.

### Mouse Xenografts

SW1088 cells (transduced with EV or ATAD1-FLAG) were grown under normal culture conditions, as described above. Cells (3 × 10^6^) were mixed 1:1 with Matrigel (Corning®) and injected into one flank per mouse. Mice were male NOD/SCID aged 13-15 weeks. Tumor volumes were monitored biweekly using a Biopticon TumorImager. Animal experiments were conducted in accordance with The University of Utah IACUC.

### Patient data

Outcome data from patients with metastatic, castration-resistant prostate cancer (mCRPC) were downloaded from TCGA via cBioPortal. Patients were stratified into three groups based on status of *ATAD1* and *PTEN* (unaltered vs. null). Raw data are available as a table in supplemental files.

### Bacterial transformation

For cloning, *E. coli* DH5α competent cells (New England Biolabs) were transformed according to the manual provided by the manufacturer and grown on LB agar plates at 37° C overnight. For cloning of lentiviral and retroviral vectors, NEB Stable competent cells were used (NEB C3040I).

### E. cloni cells

For cloning, E cloni10G competent cells were transformed according to the manual provided by the manufacturer (Lucigen) and grown on LB agar plates at 37° C overnight.

### BL21-DE3 pRIL cells

For protein expression, *E. coli* BL21(DE3) containing a pRIL plasmid and a protein expression vector were grown in terrific broth at 37° C until an OD_600_ of 0.6-1.0. Cultures were induced with isopropyl-1-thio-β-D-galactopyranoside (IPTG) at a final concentration of 1 mM and grown at room temperature for an additional 3-4 h.

### Production of soluble constructs

#### Δ1-32 Msp1 and Δ1-39ATAD1

The gene encoding the soluble region of *S. cerevisiae* Msp1 (Δ1-32) was PCR amplified from genomic DNA and subcloned into a pET28a derivative (Novagen) encoding an N-terminal 6xHis tag followed by a TEV protease cleavage site. The soluble region of *Rattus norvegicus* ATAD1 (Δ1-39) was PCR amplified from a plasmid containing ATAD1 cDNA (GE Healthcare). All insertions and deletions were performed by standard PCR techniques. Site-specific mutagenesis was carried out by QuickChange PCR. All constructs were verified by DNA sequencing.

Plasmids encoding soluble Msp1, ATAD1, or their mutants were purified as described previously (Wohlever et al., 2017). Plasmids were transformed into *E. coli* BL21(DE3) containing a pRIL plasmid and expressed in terrific broth at 37° C until an OD_600_ of 0.6-1.0, cultures were induced with 1 mM IPTG and grown at room temperature for an additional 3-4 h. Cells were harvested by centrifugation, and resuspended in Msp1 Lysis Buffer (20 mM Tris pH 7.5, 200 mM KAc, 20 mM Imidazole, 0.01 mM EDTA, 1 mM DTT) supplemented with 0.05 mg/mL lysozyme (Sigma), 1 mM phenylmethanesulfonyl fluoride (PMSF) and 500 U of universal nuclease (Pierce), and lysed by sonication. The supernatant was isolated by centrifugation for 30 min at 4° C at 18,500 x g and purified by Ni-NTA affinity chromatography (Pierce) on a gravity column. Ni-NTA resin was washed with 10 column volumes (CV) of Msp1 Lysis Buffer and then 10 CV of Wash Buffer (Msp1 Lysis buffer with 30 mM Imidazole) before elution with Lysis Buffer supplemented with 250 mM imidazole. Purification of soluble ATAD1 also included the addition of ATP to stabilize the protein. ATP was added to a final concentration of 2 mM after sonication and again after elution from the nickel resin.

The protein was further purified by size exclusion chromatography (SEC) (Superdex 200 Increase 10/300 GL, GE Healthcare) in 20 mM Tris pH 7.5, 200 mM KAc, 1 mM DTT. Peak fractions were pooled, concentrated to 5-15 mg/ml in a 30 kDa MWCO Amicon Ultra centrifugal filter (Pierce) and aliquots were flash-frozen in liquid nitrogen and stored at -80 °C. Protein concentrations were determined by A_280_ using a calculated extinction coefficient (Expasy).

#### GST-SGTA and GST-calmodulin

GST tagged SGTA was expressed and purified as described previously (Mateja et al., 2015). The original calmodulin plasmid was a kind gift of the Hegde lab (Shao and Hegde, 2011). Calmodulin was cloned into pGEX6p1 plasmid by standard methods. GST-SGTA and GST-calmodulin were expressed as described above for soluble Msp1 constructs. Cells were harvested by centrifugation and resuspended in SGTA Lysis Buffer (50 mM Hepes pH 7.5, 150 mM NaCl, 0.01 mM EDTA, 1 mM DTT, 10% glycerol) supplemented with 0.05 mg/mL lysozyme (Sigma), 1 mM PMSF and 500 U of universal nuclease (Pierce), and lysed by sonication. The supernatant was isolated by centrifugation for 30 min at 4° C at 18,500 x g and purified by Glutathione affinity chromatography (Thermo Fisher) on a gravity column. Resin was washed with 20 column volumes (CV) of SGTA Lysis Buffer and then eluted with 3 CV of SGTA Lysis Buffer supplemented with 10 mM reduced glutathione. The protein was further purified by size exclusion chromatography (SEC) (Superdex 200 Increase 10/300 GL, GE Healthcare) in 20 Mm Tris pH 7.5, 100 mM NaCl, 0.1 mM TCEP. Peak fractions were pooled, concentrated to 10 mg/ml in a 30 kDa MWCO Spin Concentrator (Pierce) and aliquots were flash-frozen in liquid nitrogen and stored at -80 °C. Protein concentrations were determined by A_280_ using a calculated extinction coefficient (Expasy).

### Production of membrane proteins

#### BIM and Fis1

*Homo sapiens* BimL or *S. cerevisiae* Fis1 TMD +/- 5 flanking amino acids (residues 126-155) was cloned in place of the Sec22 TMD in the SumoTMD construct described previously (Wang et al., 2010; Wohlever et al., 2017). These constructs have N-terminal His_6_ and 3x Flag tags and a C-terminal opsin glycosylation site (11 residues). A 3C protease site was added immediately after the His tag by standard PCR methods. The resulting constructs are His_6_-3C-3xFlag-Sumo-thrombin-BimL-Opsin and His_6_-3C-3xFlag-Sumo-thrombin-Fis1(126-155)-Opsin.

Expression plasmids for SumoTMD were transformed into *E. coli* BL21(DE3) containing a pRIL plasmid and expressed in terrific broth at 37° C until an OD_600_ of 0.6-0.8, cultures were induced with 0.4 mM IPTG and grown at 20° C for an additional 3-4 h. Cells were harvested by centrifugation, and resuspended in SumoTMD Lysis Buffer (50 mM Tris pH 7.5, 300 mM NaCl, 10 mM MgCl_2_, 10 mM Imidazole, 10% glycerol) supplemented with 0.05 mg/mL lysozyme (Sigma), 1 mM PMSF and 500 U of universal nuclease (Pierce), and lysed by sonication.

Membrane proteins were solubilized by addition of n-dodecyl-β-D-maltoside (DDM) to a final concentration of 1% and rocked at 4° C for 30’. Lysate was cleared by centrifugation for at 4° C for 1 h at 35,000 x g and purified by Ni-NTA affinity chromatography.

Ni-NTA resin was washed with 10 column volumes (CV) of SumoTMD Wash Buffer 1 (50 mM Tris pH 7.5, 500 mM NaCl, 10 mM MgCl_2_, 10 mM imidazole, 5 mM β-mercaptoethanol (BME), 10% glycerol, 0.1% DDM). Resin was then washed with 10 CV of SumoTMD Wash Buffer 2 (same as Wash Buffer 1 except with 300 mM NaCl and 25 mM imidazole) and 10 CV of SumoTMD Wash Buffer 3 (same as Wash Buffer 1 with 150 mM NaCl and 50 mM imidazole) and then eluted with 3 CV of SumoTMD Elution Buffer (same as Wash Buffer 3 except with 250 mM imidazole).

The protein was further purified by size exclusion chromatography (SEC) (Superdex 200 Increase 10/300 GL, GE Healthcare) in 50 mM Tris pH 7.5, 150 mM NaCl, 10 mM MgCl_2_, 5 mM BME, 10% glycerol, 0.1% DDM. Peak fractions were pooled and concentrated in a 30 kDa MWCO spin concentrator (Pierce). Sample was then incubated with 3C Protease at a 1:100 ratio at 4° C overnight to remove the His tag. The following day, the sample was run over Ni-NTA resin equilibrated in Lysis Buffer to remove 3C protease, His tag, and uncleaved proteins. Flow through was collected, aliquoted, and flash-frozen in liquid nitrogen and stored at -80 °C. Protein concentrations were determined by A_280_ using a calculated extinction coefficient (Expasy).

#### Msp1

Full-length *S. cerevisiae* Msp1 was PCR amplified from genomic DNA, subcloned into a pET21b derivative with a C-terminal 6xHis tag and expressed as described above for the soluble constructs. Cells were lysed by sonication and the insoluble fraction was harvested by centrifugation for 1 h at 4° C at 140,000 x g. After resolubilizing for 16 h in Msp1 Lysis Buffer containing 1% DDM (Bioworld), the detergent-soluble supernatant was isolated by centrifugation for 45 min at 142,000 x g and purified by Ni-NTA affinity chromatrography and SEC as described above for the soluble constructs, except that all buffers contained 0.05% DDM. Peak fractions were concentrated in 100 kDa MWCO Amicon Ultra centrifugal filter (Millipore). Protein concentrations were determined by A_280_ using a calculated extinction coefficient (Expasy) and aliquots were flash frozen in liquid nitrogen.

### Reconstitution of Msp1 activity in proteoliposomes

#### Liposome preparation

Liposomes mimicking the lipid composition of the yeast outer mitochondrial membrane were prepared as described (Kale et al., 2014). Briefly, a 25 mg lipid film was prepared by mixing chloroform stocks of chicken egg phosphatidyl choline (Avanti 840051C), chicken egg phosphatidyl ethanolamine (Avanti 840021C), bovine liver phosphatidyl inositol (Avanti 840042C), synthetic DOPS (Avanti 840035C), and synthetic TOCL (Avanti 710335C) at a 48:28:10:10:4 molar ratio with 1 mg of DTT. Nickel liposomes were made as described above, except, 1,2-dioleoyl-sn-glycero-3-[N-(5-amino-1-carboxypentyl)iminodiacetic acid)succinyl] Nickel salt (Avanti 790404) was used at a molar ratio of 2% and DOPS was dropped from 10% to 8%.

Chloroform was evaporated under a gentle steam of nitrogen and then left on a vacuum (<1 mTorr) overnight. Lipid film was resuspended in Liposome Buffer (50 mM Hepes KOH pH 7.5, 15% glycerol, 1 mM DTT) to a final concentration of 20 mg/mL and then subjected to five freeze-thaw cycles with liquid nitrogen. Liposomes were extruded 15 times through a 200 nm filter at 60° C, distributed into single use aliquots, and flash frozen in liquid nitrogen.

#### Proteoliposome preparation

For extraction assays with full-length Msp1, proteoliposomes were prepared by mixing 1 µM Msp1, 1 µM TA protein (SumoTMD), and 2 mg/mL of mitochondrial liposomes in Reconstitution Buffer (50 mM Hepes KOH pH 7.5, 200 mM potassium acetate, 7 mM magnesium acetate, 2 mM DTT, 10% sucrose, 0.01% sodium azide, and 0.1% deoxy big chaps)(Zhang et al., 2013). For extraction assays with soluble Msp1/ATAD1, proteoliposomes were prepared by mixing 1 µM TA protein (SumoTMD), and 2 mg/mL of Nickel liposomes in Reconstitution Buffer. Detergent was removed by adding 25 mg of biobeads and rotating the samples for 16 h at 4° C. After removing biobeads, unincorporated TA protein was pre-cleared by incubating the reconstituted material with excess (5 µM) GST-SGTA and GST-Calmodulin and passing over a glutathione spin column (Pierce #16103); the flow through was collected and used immediately for dislocation assays.

### Extraction Assay

Extraction assays contained 60 µL of pre-cleared proteoliposomes, 5 µM GST-SGTA, 5 µM calmodulin, and 2 mM ATP and the final volume was adjusted to 200 µL with Extraction Buffer (50 mM Hepes KOH pH 7.5, 200 mM potassium acetate, 7 mM magnesium acetate, 2 mM DTT, 0.1 µM calcium chloride). Samples were incubated at 30° C for 35 min and then loaded onto a glutathione spin column. Columns were washed 4x with Extraction Buffer and eluted with the same buffer supplemented with 20 mM glutathione pH 8.5. Samples were loaded onto stain free gels, imaged, and then transferred to a PVDF membrane and blotted as indicated in the key resource table. To account for variability in reconstitution efficiency and western blotting, a new reconstitution and dislocation assay with wild-type Msp1 was done in parallel with each mutant Msp1. Figures are representative of N > 3 separate reconstitutions. Note that the “input” lane is diluted 5x relative to the “elution” lane.

### Quantification and Statistical Analysis

To account for variability in reconstitution efficiency and western blotting, a new reconstitution and dislocation assay with wild-type Msp1 was done in parallel with each Msp1 mutant. Figures are representative of N > 3 separate reconstitutions. Dislocation efficiency was quantified by comparing the amount TA protein in the “elution” lane with the amount of substrate in the “input” lane.

## KEY RESOURCES TABLE

| REAGENT or RESOURCE | SOURCE | IDENTIFIER |
| --- | --- | --- |
| Antibodies |  |  |
| Anti-Flag | Sigma-Aldrich | #F7425,<br>RRID:AB_439687 |
| Anti-V5 | Abcam | #ab9116,<br>RRID:AB_307024 |
| Anti-GFP | Cell Signaling | #2956S |
| Anti-ATAD1/Thorase | NeuroMab | 75-157 |
| Anti-PTEN | Cell Signaling | #9188 |
| Anti-beta Actin | Cell Signaling | #4970 |
| Anti-alpha tubulin | Cell Signaling | #3873 |
| Anti-BIM | Cell Signaling | #2933 |
| Anti-BAK | Cell Signaling | #12105 |
| Anti-Ubiquitin, rabbit | Cell Signaling | #43124 |
| Anti-Ubiquitin, mouse | Abcam | #ab7254 |
| Anti-GAPDH | Cell Signaling | #97166S |
| Anti-phospho p44/42 MAPK (ERK1/2) | Cell Signaling | #4370 |
| Anti-p44/42 MAPK (ERK1/2) | Cell Signaling | #9101 |
| Anti-PARP | Cell Signaling | #9532 |
| Anti-Caspase 3 | Cell Signaling | #14220S |
| Goat Anti-Mouse IgG (H&L) Antibody Dylight™<br>800 Conjugated | Rockland | #610-145-002-0.5,<br>RRID: AB_10703265 |
| Donkey anti-Rabbit IgG (H+L) Highly Cross-Adsorbed Secondary Antibody, Alexa Fluor 680 | Invitrogen | #A10043,<br>RRID: AB_2534018 |
| Goat Anti-Mouse IgG (H+L) Antibody, Alexa<br>Fluor 680 Conjugated | Invitrogen | #A21057,<br>RRID:AB_141436 |
| Donkey Anti-Rabbit IgG (H&L) Antibody<br>Dylight™ 800 Conjugated | Rockland | #611-145-002-0.5,<br>AB_11183542 |
| Goat anti-Mouse IgG (H+L), Superclonal™<br>Recombinant Secondary Antibody, HRP | Thermo Fisher<br>Scientific | #A28177,<br>RRID:AB_2536163 |
| Goat anti-Rabbit IgG (H+L), HRP | Proteintech | RRID:AB_2722564 |
| Bacterial and Virus Strains |  |  |
| Biological Samples |  |  |
| Chemicals, Peptides, and Recombinant Proteins |  |  |
| DDM | GoldBio | DDM25 |
| Lipofectamine 3000 | ThermoFisher | L3000008 |
| Lipofectamine RNAiMAX | Invitrogen | 13778150 |
| MitoTracker Red CMXRos | Invitrogen | M7512 |
| Crystal Violet | Sigma | C0775 |
| RIPA Buffer | Cell Signaling | 9806 |
| SE Cell Line 4D-Nucleofector™ X Kit L | Lonza | V4XC-1012 |
| Adenosine Triphosphate | Acros Organics | AC10280-0100 |
| bovine liver phosphatidyl inositol | Avanti | 840042C-10mg |
| Synthetic DOPS | Avanti | 840035C-10mg |
| Synthetic DOGS-Ni-NTA | Avanti | 790404C-5mg |
| Chicken egg phosphatidyl ethanolamine | Avanti | 840021C-25mg |
| Chicken egg phosphatidyl choline | Avanti | 840051C-200mg |
| Synthetic TOCL | Avanti | 710335C-25mg |
| Critical Commercial Assays |  |  |
| Pierce BCA Assay | Thermo | 23225 |
| Cell Titer Glo Luminescent Viability Assay | Promega | G7572 |
| Deposited Data |  |  |
| Experimental Models: Cell Lines |  |  |
| HEK293T cells | ATCC | #CRL-11268,<br>RRID:CVCL_192<br>6 |
| Jurkat E6.1 | ATCC | TIB-152 |
| H4 | ATCC | HTB-148 |
| RPMI-7951 | ATCC | HTB-66 |
| SW1088 | ATCC | HTB-12 |
| PC3 | ATCC | CRL-1435 |
| HGC27 | Sigma Aldrich | 94042256 |
| Experimental Models: Organisms/Strains |  |  |
| Oligonucleotides |  |  |
| Recombinant DNA |  |  |
| psPAX2 | Addgene | #12259,<br>RRID:Addgene_12<br>259 |
| pMD2.G | Addgene | #12260,<br>RRID:Addgene_12<br>260 |
| pSpCas9(BB)-2A-GFP (PX458) | Addgene | 48138 |
| LentiCRISPRv2GFP | Addgene | 82416 |
| pLenti-Blast | Addgene | 17486 |
| PQCXIP | Clontech |  |
| pLVX-TetOne-Puro | Takara | 631844 |
| pET21: Msp1-His (*S. cerevisiae*) | Wohlever et al. 2017 |  |
| pET28: His-TEV-□1-32-Msp1 (*S. cerevisiae*) | Wohlever et al. 2017 |  |
| pET28: His-TEV-□1-39-ATAD1 (*R. norvegicus*) | This study |  |
| pET28: His-Flag-Sumo-Sec22 TMD-Op sin | Wang et al. 2010 |  |
| pET28: His-Flag-Sumo-BimL-Op sin | This study |  |
| pET28: His-Flag-Sumo-Fis1 TMD-Op sin | This study |  |
| pGEX6p1: GST-SGTA | Mateja et al. 2015 |  |
| pGEX6p1: GST-Calmodulin | Shao & Hegde 2011 |  |
| Software and Algorithms |  |  |
| metap | Michael Dewey,<br>2020 | http://www.dewey.myzen.co.uk/meta/meta.html |
| R | R Core Team | https://www.r-project.org |
| ggplot2 | Wickham, 2009 | http://ggplot2.org |
| Other |  |  |

