## Supplemental Figures for "Co-deletion of *ATAD1* with *PTEN* primes cells for BIM-mediated apoptosis"

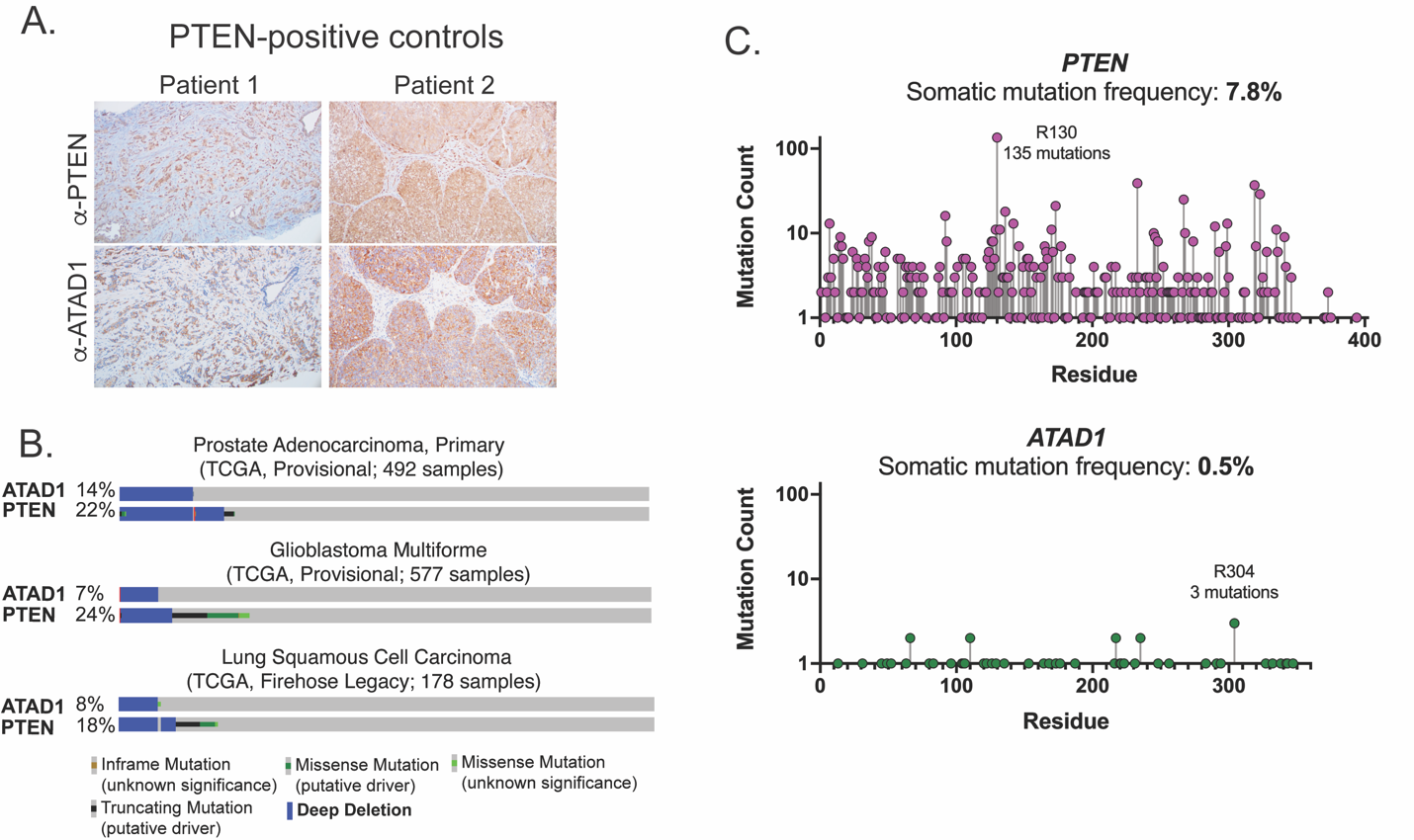

Extended Data Figure 1: ATAD1 is co-deleted with PTEN as a passenger. (A) Representative histology of tumor samples from patients with PTEN-positive tumors, as a control for the PTEN-negative tumors shown in Figure 1 (B) Oncoprint of alterations at the ATAD1 and PTEN loci in three different TCGA studies (C) Somatic mutations in the *PTEN* or *ATAD1* loci, from TCGA Pan-Cancer Atlas studies (32 studies; n = 10,528 samples); note log­arithmic scale on y-axis.

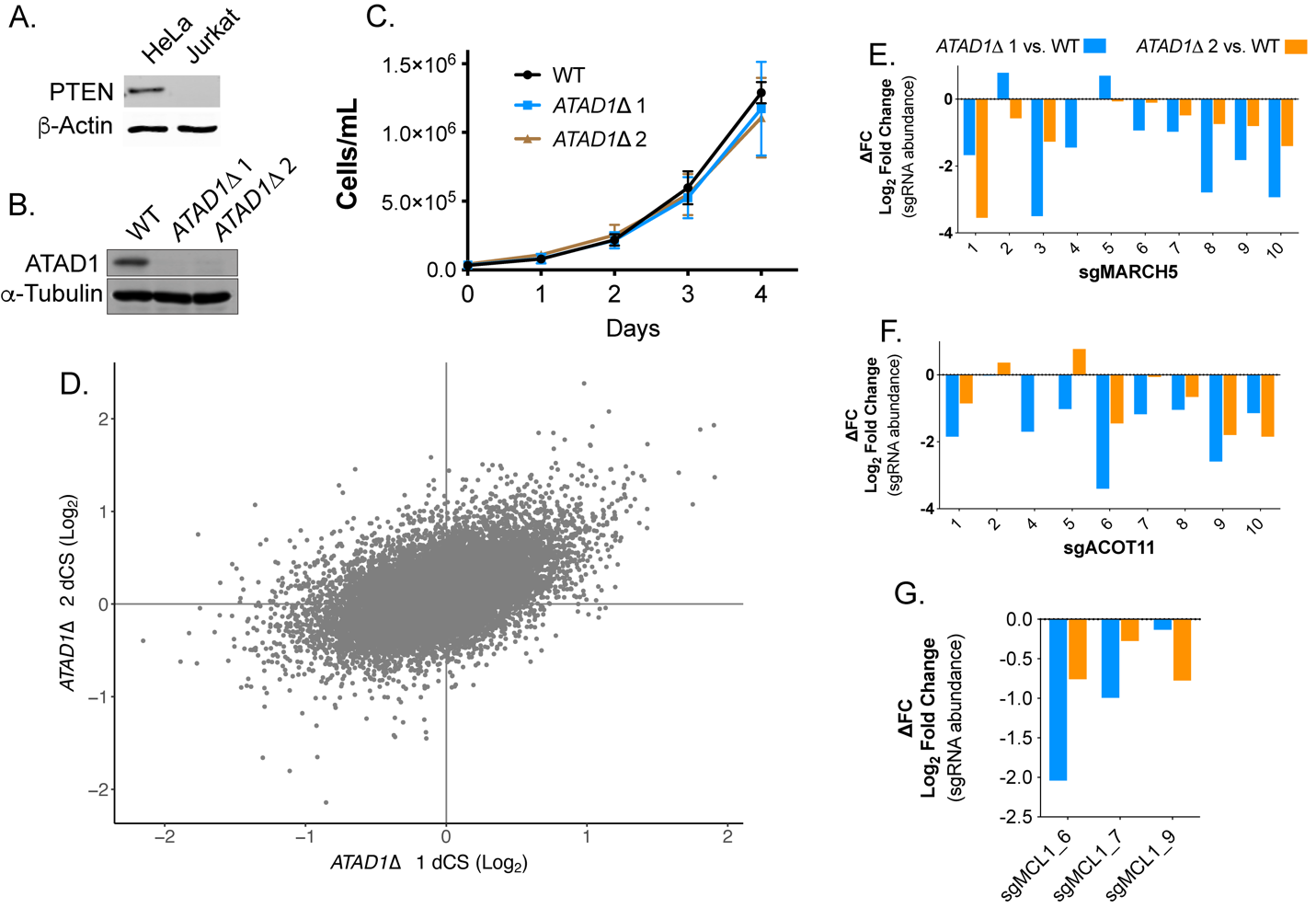

Extended Data Figure 2: Supporting data for CRISPR screens. (A) Western blot verification of PTEN-deficiency of Jurkat cells (B) Western blot verification of ATAD1-deficiency of *ATAD1*∆ cell lines (C) Proliferation of WT and *ATAD1*∆ cell lines over 4 days; mean ± SD for n = 3 independent experiments (D) Differential CRISPR scores for the two *ATAD1*∆ clonal cell lines relative to WT; Pearson coefficient = 0.51, P = 2.16 x 10^-16^ (E) Differential CRISPR scores by sgRNA for *MARCH5* and (F) *ACOT11* (G) dCS values for three sgRNAs that were represented in the initial time point (after puromycin selection, before outgrowth) targeting *MCL1*

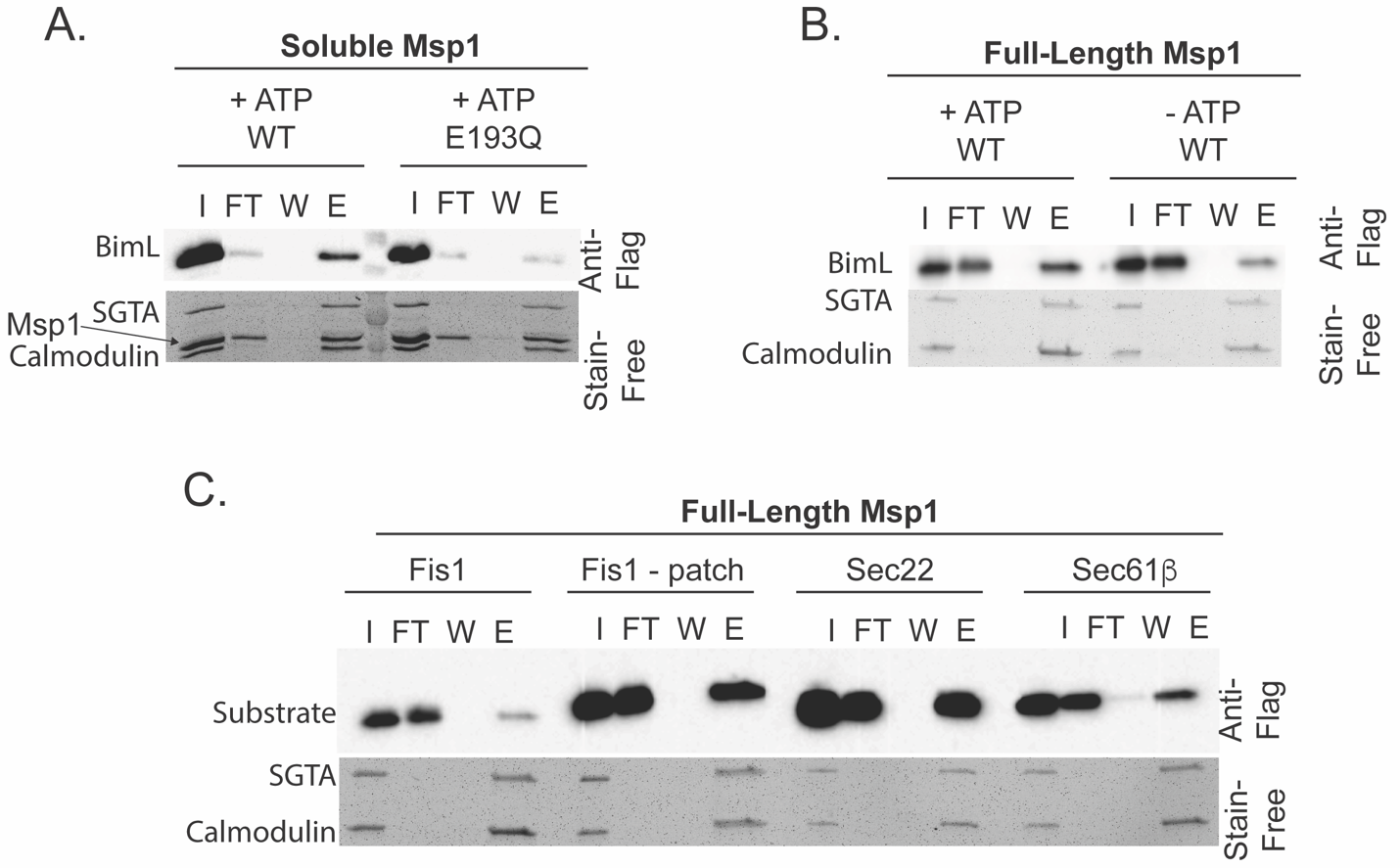

Extended Data Figure 3: Validation of proteoliposome extraction assay (A) Soluble His-Msp1 and full-length Msp1 (B) extract BIM_L_ from proteoliposomes. (C) The extraction assay recapitulates physiologic substrate selectivity of Msp1. Fis1 is extracted when a known Msp1 recognition motif consisting of a hydrophobic patch of residues from Pex15, is inserted N terminal to the TMD (“Fis1 - patch”). Sec22 and Sec61b are positive controls demonstrating that Msp1 can recognize ER-native TA proteins.

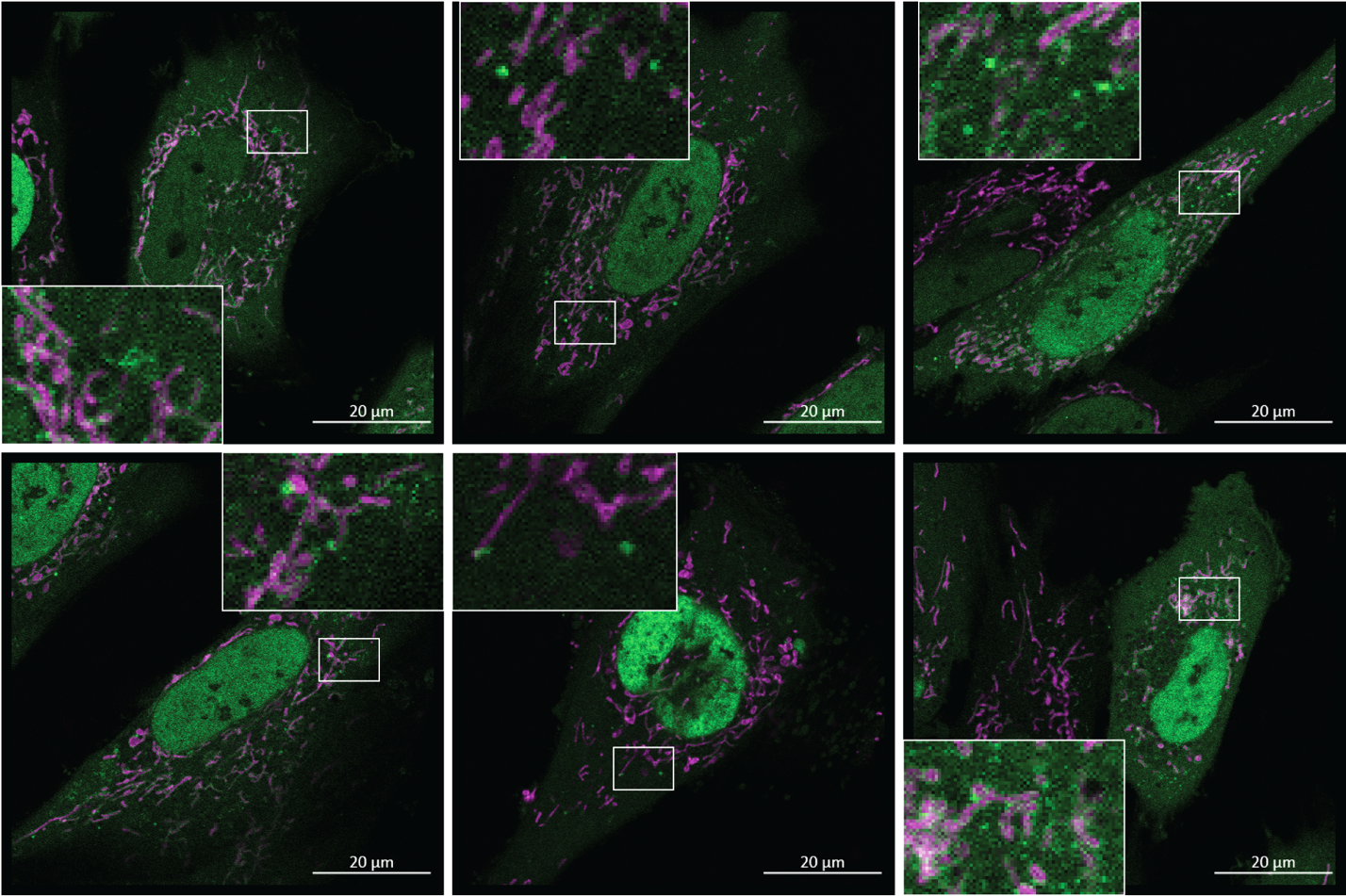

Extended Data Figure 4: Examples of GFP-positive puncta induced by bortezomib treatment in SW1088 cells expressing ATAD1 and GFP-BIM_EL_∆BH3. Images were taken by a blinded investigator and are representative of 3 independent experiments.

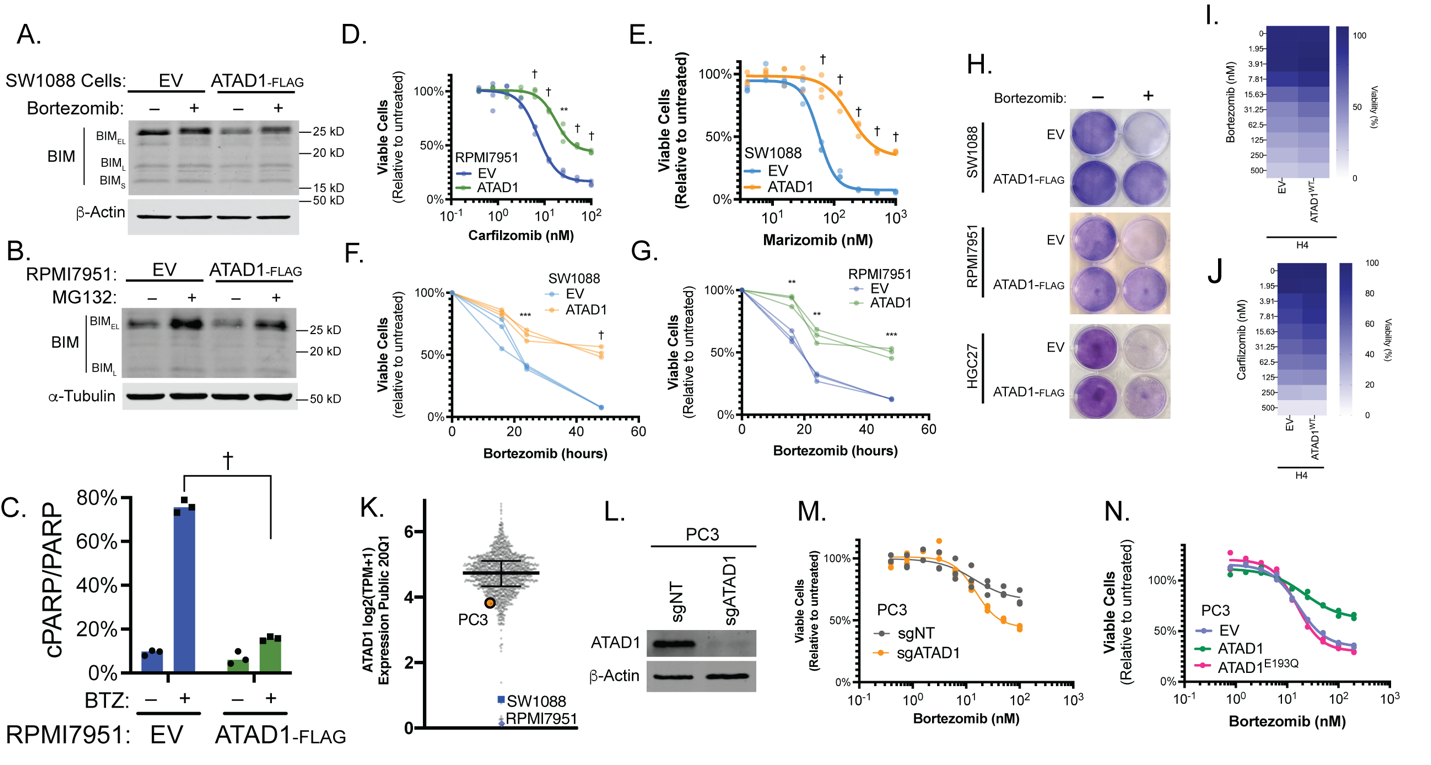

Extended Data Figure 5: Additional data on ATAD1 status and proteasome inhibition. (A) Western blot of SW1088 cells treated with bortezomib or RPMI7951 cells treated with MG132 (B) and assessed for BIM protein levels (C) Quantification of PARP cleavage in RPMI7951 cells treated with 10 nM bortezomib for 8 hr. Values and median (bar) from three independent experiments are shown. (D) Viability of RPMI7951 cells treated with carfilzomib for 16 hr, n = 3 biological replicates, 2 independent experiments (E) Viability of SW1088 cells treated with marizomib for 24 hr, n = 3 biological replicates, 2 independent experiments (F) Viability of SW1088 cells treated with bortezomib (3.9 nM) for the indicated durations, n = 3 biological replicates (G) Viability of RPMI7951 cells treated with bortezomib (3.1 nM) for the indicated durations, n = 3 biological replicates (H) SW1088 cells were treated with 50 nM bortezomib for 48 hr and RPMI7951 and HGC27 cells for 24 hr and then stained with crystal violet, n = 2 independent experiments each. H4 cells were treated with bortezomib (I) or carfilzomib (J) for 24 hr and viability assessed by CellTiterGlo; mean of 2 biological replicates is shown (K) Expression of ATAD1 in DepMap cell lines, highlighting PC3 cells (ATAD1 hemizygous), and two Del10q23 cell lines (L) Generation of ATAD1-deficient, polyclonal PC3 cells using stable expression of sgATAD1 (in LCv2G), with non-targeting sgRNA as a control (sgNT) (M) PC3 cells were transduced with EV, ATAD1^WT^-FLAG/HA, or ATAD1^E193Q^-FLAG/HA and treated with bortezomib for 24 hr. (N) PC3 cells as shown in (G) were treated with bortezomib overnight and viability assessed by CellTiterGlo. PC3 cell experiments are representative of two independent experiments with two (N) or three (M) biological replicates.
